# Widespread natural selection on metabolite levels in humans

**DOI:** 10.1101/2023.02.07.527420

**Authors:** Yanina Timasheva, Kaido Lepik, Orsolya Liska, Balázs Papp, Zoltán Kutalik

## Abstract

Natural selection acts ubiquitously on complex human traits, predominantly constraining the occurrence of extreme phenotypes (stabilizing selection). These constrains propagate to DNA sequence variants associated with traits under selection. The genetic imprints of such evolutionary events can thus be detected via combining effect size estimates from genetic association studies and the corresponding allele frequencies. While this approach has been successfully applied to high-level traits, the prevalence and mode of selection acting on molecular traits remains poorly understood. Here, we estimate the action of natural selection on genetic variants associated with metabolite levels, an important layer of molecular traits. By leveraging summary statistics of published genome-wide association studies with large sample sizes, we find strong evidence of stabilizing selection for 15 out of 97 plasma metabolites, with an overrepresentation of amino acids among such cases. Mendelian randomization analysis revealed that metabolites under stronger stabilizing selection display larger effects on key cardiometabolic traits, suggesting that maintaining a healthy cardiometabolic profile may be an important source of selective constraints on the metabolome. Metabolites under strong stabilizing selection in humans are also more conserved in their concentrations among diverse mammalian species, suggesting shared selective forces across micro and macroevolutionary time scales. Finally, we also found evidence for both disruptive and directional selection on specific lipid metabolites, potentially indicating ongoing evolutionary adaptation in humans. Overall, this study demonstrates that variation in metabolite levels among humans is frequently shaped by natural selection and this may be acting indirectly through maintaining cardiometabolic fitness.

## Introduction

Human metabolites provide a unique insight into metabolic pathways underlying health and disease, and can serve as a useful tool for precision medicine with multiple applications, including discovery of new therapeutic targets and development of novel protocols for diagnostics or monitoring the progression of the disease and the efficacy of treatment ^1^. Recent advances in metabolomics research have identified a number of biochemical processes involved in the pathogenesis of complex diseases, such as cancer, atherosclerosis, and diabetes ^2^.

Intermediate metabolites can also help to elucidate the influence of natural selection due to the evolutionary advantages and disadvantages resulting from the ability of living organisms to produce compounds with functions beneficial or detrimental for fitness. While dysregulation of several specific metabolites has been linked to human diseases, potentially indicating strong stabilizing selection to preserve their levels, the direction and strength of natural selection shaping metabolite levels is generally unknown. Several lines of observations suggest that stabilizing selection on metabolite levels might be prevalent. First, evolutionary distant species show substantial similarities in metabolite levels, indicating widespread evolutionary conservation of the metabolome ^3; 4^. Second, the levels of central metabolites obey simple optimality principles, indicating that metabolite levels might represent optimal values ^5^. However, not all metabolites are expected to be under equally strong stabilizing selection and there might be larger room for selectively neutral alterations for some metabolites than for others. Indeed, a recent multi-species comparison revealed wide differences in the extent of conservation of individual metabolite concentrations during evolution, likely driven by differing amounts of functional constraints across metabolites ^6^. Furthermore, a remarkable acceleration of metabolome evolution has been reported in the human lineage compared to other primates, potentially indicating the action of directional selection on specific metabolites ^7^. However, the general patterns of selection shaping human metabolite concentrations remain essentially unknown.

Evidence for stabilizing selection acting on a particular trait can be inferred from the relationship between the multivariable effect size (*b*) and minor allele frequency (MAF) of genetic variants responsible for its regulation ^8^. The observed omnigenic architecture of complex traits suggests that a large number of trait-associated genetic variants have very small effect, while only a few of them have larger effects. Individuals carrying alleles associated with larger (detrimental) effects on a trait under strong (stabilizing) selection will have higher chance to have decreased fitness and hence will tend to be reduced/purged from the gene pool. This results in a decreased allele frequency. It is generally viewed that most traits under (stabilizing) negative selection have an optimal value for fitness and individuals with larger deviations in either direction tend to be selected out with increasing probability. Hence, it is reasonable to assume that in such a case there is an inverse relationship between the squared effect size and MAF (or the variability of the genotype, i.e. 2*MAF*(1-MAF)). Stronger selection leads to the sharper decline in MAF upon increase in effect size. Therefore, it has been proposed to estimate selection strength acting on a phenotype, denoted by α, as the value that best fits the b^2^ ∼ [2*MAF*(1-MAF)]^α^ relationship for the given phenotype ^9^.

More importantly, simulation studies demonstrated that negative *α* values point to stabilizing, while positive *α* values to either directional or disruptive selection ^8^. For typical complex traits a reasonable *α* was estimated to be around -0.25 ^10^. It was shown that stratifying heritability models for functional annotations, LD scores and MAF can improve heritability estimation and trait prediction^11^. Hence, identifying the signatures of negative selection is important for understanding both the genetic underpinnings of phenotypic variation and evolution.

Such methodology has not yet been applied to molecular traits, such as metabolite concentrations, due to the lack of statistical power because of the unavailability of sufficiently large sample size. The recent emergence of genome-wide association summary statistics for metabolites^12^ provided the first such opportunity. To leverage the newly available data, we reliably estimated selection strength for 97 out of 135 metabolites from seven biochemical classes. First, we compared the performances of weighted and unweighted LDAK-Alpha and BLD-LDAK-Alpha models using summary statistics data for 59 complex traits available from the UK Biobank. We then applied the weighted model to obtain the selection strength signatures for 135 metabolites using the summary statistics data generated in a cross-platform meta-analysis of genetic effects on levels of blood metabolites measured in large-scale population-based studies ^12^. Next, we explored the imprint of directional selection for the studied metabolites by analysing the relationship between effect allele frequency (EAF) and the effect sizes of lead SNPs associated with the studied metabolites at suggestively significant (p < 5 × 10^−6^) or at a genome-wide significant level (p < 5 × 10^−8^). We also investigated the causal relationship between metabolites and six clinically important complex traits using Mendelian randomization (MR) in order to explore the relationship between the selection strength estimates for the studied metabolites and their impact on cardiometabolic traits contributing to fitness. Finally, we used an orthogonal measure of evolutionary constraints by comparing cross-species metabolite concentrations among mammals, and compared it with the strength for stabilizing selection obtained from human GWAS data.

## Materials and methods

We analysed genome-wide summary statistics for 135 metabolites, including amino acids, biogenic amines, acylcarnitines, lysophosphatidylcholines, phosphatidylcholines, sphingomyelins, and hexoses, made available recently from a large (up to 86,507 individuals) genome-wide meta-analysis study ^12^.

### Stabilizing selection

We utilised the SumHer approach implemented in the LDAK software ^13^, having compared the performance of different heritability models to identify the optimal version for assessing the selection strength. The tested models were based on the LDAK model where expected per-SNP-heritability of a SNP depends on both the LD and MAF:

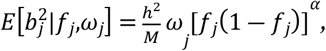

where 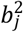is the multivariable squared effect size of SNP *j, f*_*j*_ is the observed MAF, *h*^2^ is the trait heritability, *M* is the number of causal markers, *ω*_*j*_ represents the SNP weighs, which are inversely proportional to the LD score (computed based on the local LD structure), while parameter *α* represents the strength of stabilizing selection. This parameter is estimated from the data via profile likelihood maximization. In brief, we fixed the selection strength parameter (*α*) and maximized the likelihood function for *h*^2^. This procedure was repeated for 31 values of *α*, ranging from -1 to 0.5 with step size of 0.05. Finally, we plotted the maximum likelihood values against the value of *α* and fitted a quadratic polynomial to these points. Once the estimates for the coefficients of the polynomial were established, the maximum and the negative Hessian (of its second derivative with respect to *α*) were computed, yielding the maximum likelihood estimator for *α* and its variance. For some traits, we observed that the optimal *α* values fall outside of the [-1, 0.5] range and in such situations, we extended the range to either [-2.0, 1.0] or [-1.0, 4.0], as necessary.

We adjusted this procedure to use more complex, stratified heritability estimation with 65 genomic annotations (resulting in stratum-specific heritability estimates). This BLD-LDAK-Alpha model extends the LDAK-Alpha model by adding 65 annotations provided by Hujoel et al ^14^. We tried simpler models (without annotation, or equal weights (*ω*_*j*_ =1)) and compared the performance of these models using the summary statistics data for 59 well-studied complex traits (including height, BMI, SBP, etc.) from the UK Biobank (N= 361,194). The estimates for *α* from the BLD-LDAK-Alpha model came closest to the ones reported in previous studies ^,8; 11^, therefore this latter model was applied to investigate the selection signatures for 135 metabolites (with the sample size varying between 9’363 and 86’507) ^12^.

### Directional selection

We have also investigated the imprint of directional selection by exploring the relationship between EAF and the effect sizes of lead SNPs associated with the studied metabolites at suggestively significant (p < 5 × 10^−6^) or at a genome-wide significant level (p < 5 × 10^−8^). The rationale behind is that if the average population level of a metabolite is lower than its optimal value, trait-increasing alleles tend to have higher frequency due to positive selection. While this analysis may be suboptimal since marginal effects are used (instead of multivariate ones), the approach could be used to detect at least the directionality of such selection.

### Mendelian randomization analysis

We applied inverse variance weighted MR approach^15^ to explore the causal relationship between metabolites and five complex traits (BMI, IHD, SBP, LDL, and T2D) using genetic variants associated with the metabolites as instrumental variables. The obtained MR causal effects were used to further interrogate the relationship between the selection strength estimates for the studied metabolites and cardiometabolic traits contributing to the decreased fitness.

### Estimated evolutionary conservation scores of metabolites in mammals

To assess the evolutionary conservation of metabolite levels in mammals, we used a previous cross-species metabolomics study which quantified the relative concentrations of >250 metabolites in four organs (brain, heart, kidney and liver) of 26 mammalian species^3^. To calculate metabolite conservation scores (i.e. the extent to which the concentration of any given metabolite is permitted to change over the course of evolution) we fit a Brownian Motion model of trait evolution on each metabolite in each organ across the phylogeny of 26 species. We defined the conservation score of each metabolite as the inverse of the rate of concentration change that was inferred from the phylogeny. In order to get a unified conservation score for each metabolite across the four organs, we first imputed all missing values (i.e. metabolite that weren’t measured in all four organs) by calculating the median conservation score of the given metabolite across the measured organs. Then, we calculated the ranks of metabolite conservation scores across the metabolome in each organ, and used the median rank value across the four organs as an aggregate measure of metabolite conservation. Only metabolites measured in more than one organ were included in this analysis, providing us with aggregate conservation scores for a total of 249 metabolites, out of which 47 were among the 97 metabolites for which selection strength could be estimated.

## Results

### Confirmation of the heritability model using UK Biobank data for complex traits

We first applied the BLD-LDAK+Alpha model implemented in the SumHer functionality of the LDAK software to 59 complex traits available in the UK Biobank (Figure S1). After testing different heritability models, we established that fitting the 65-parameter BLD-LDAK+Alpha model for the majority of the 59 of studied traits led to *α* estimates within the range of -0.9 to -0.15. Out of the 59 traits, 25 were also tested in a previous study using raw genotype data^8^. For these 25 overlapping traits we observed reasonable similarity (Figure S2) between selection strength estimates by the 65-parameter BLD-LDAK+Alpha model vs approaches based on raw genetic data: Zeng et al ^8^ (r=0.31) and Schoech et al ^9^ (r=0.39). Note that our *α* estimates showed better agreement with the two earlier estimates than the latter with each other (Figure S2).

### Metabolome-wide signatures of natural selection

Having tested the BLD-LDAK+Alpha model on complex traits, we performed the analysis to explore, for the first time, the evidence of selection for 135 metabolites for which summary statistics are available from GWAS with large sample sizes ^12^. Notably, 38 metabolites did not produce stable ML estimates for the selection parameter, most likely due to low heritability or sample size. Out of the remaining 97 metabolites (with *α* estimates ranging from -1.82 to 3.43), 66 led to selection estimates not significantly different from zero. 28 metabolites with nominally significant (*P* < 0.05) selection estimates showed stabilizing selection (*α* < 0) and 3 are estimated to be under directional or disruptive selection (*α* > 0). Figure 1 illustrates the estimated selection strength values for these metabolites. We have found strong evidence (P<0.05/97) of stabilizing selection for 15 metabolites (tyrosine, butyrylcarnitine, acetylornithine, methionine, glutamine, PC ae C38:0, glutamate, proline, PC ae C34:1, lysoPC a C16:1, PC aa C32:1, asparagine, PC aa C34:1, lysoPC a C20:3, PC ae C40:4). Evidence for the three nominally significant metabolites under disruptive / directional selection (citrulline, lyso PC a C20:4, PC ae C40:5) did not reach the adjusted statistical significance level (P < 0.05/97) (Figure S3).

**Figure 1.**
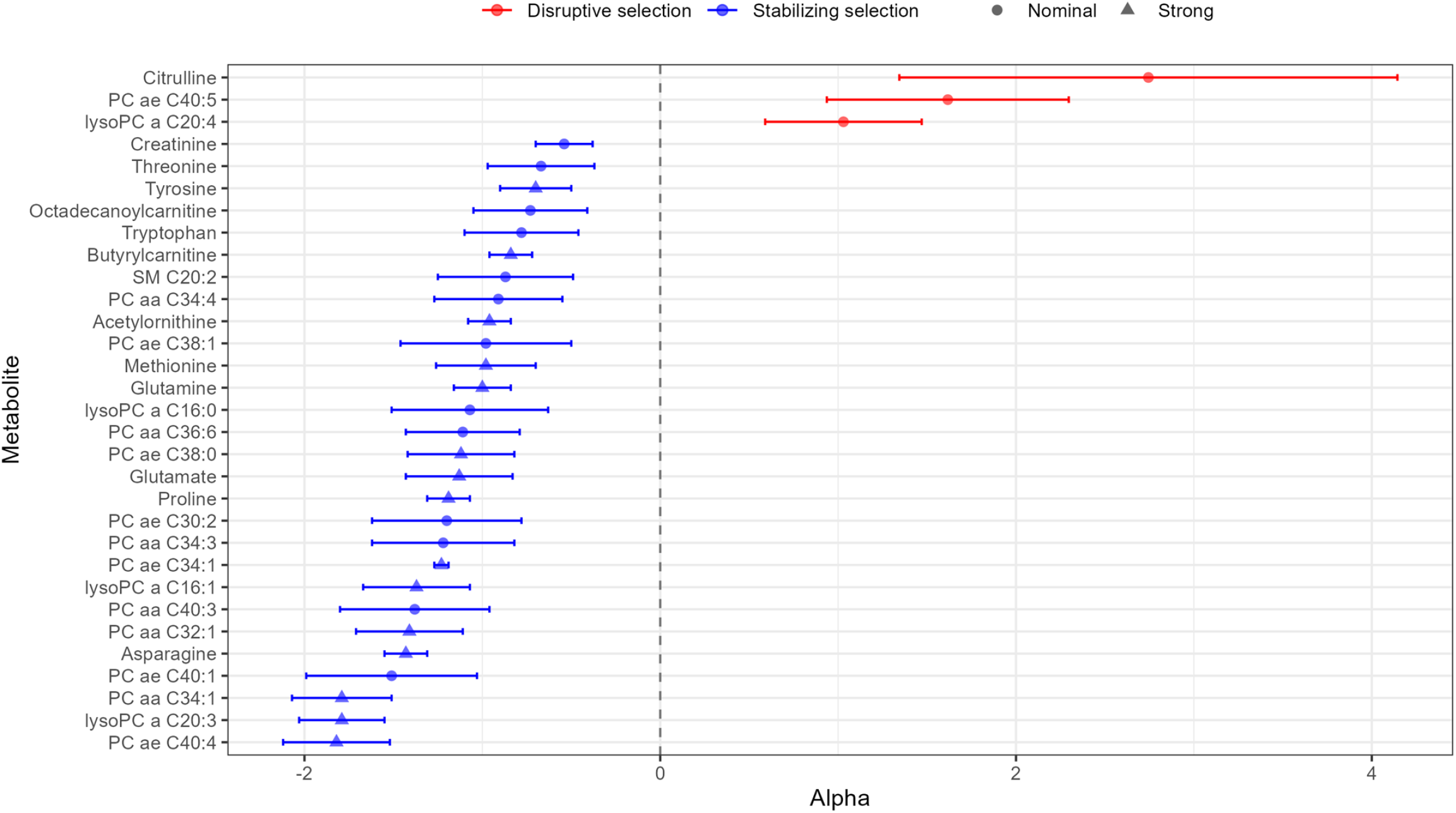
Metabolites with selection estimates showing stabilizing selection (*α* < 0) and disruptive selection (*α* > 0) at the nominally significant (*P* < 0.05) level. Circle symbols represent nominally significant selection estimates, triangles refer to metabolites with selection strength P-value surviving multiple testing correction (P<0.05/97). Error bars represent standard errors.

While metabolites showing evidence for stabilizing selection span several major compound classes, they are especially prevalent among amino acids (Figure 2). Specifically, ∼33% of amino acids and derivatives show strong evidence of stabilizing selection as compared to 10.5% of the rest of investigated metabolites (P=0.018, two-sided Fisher’s exact test). In contrast, acylcarnitines tend to have positive selection coefficients (average alpha value = 0.85, proportion of acylcarnitines with positive selection strength estimates 12/18), hinting that they may be under less stringent evolutionary constraint (either no or disruptive / directional selection).

**Figure 2.**
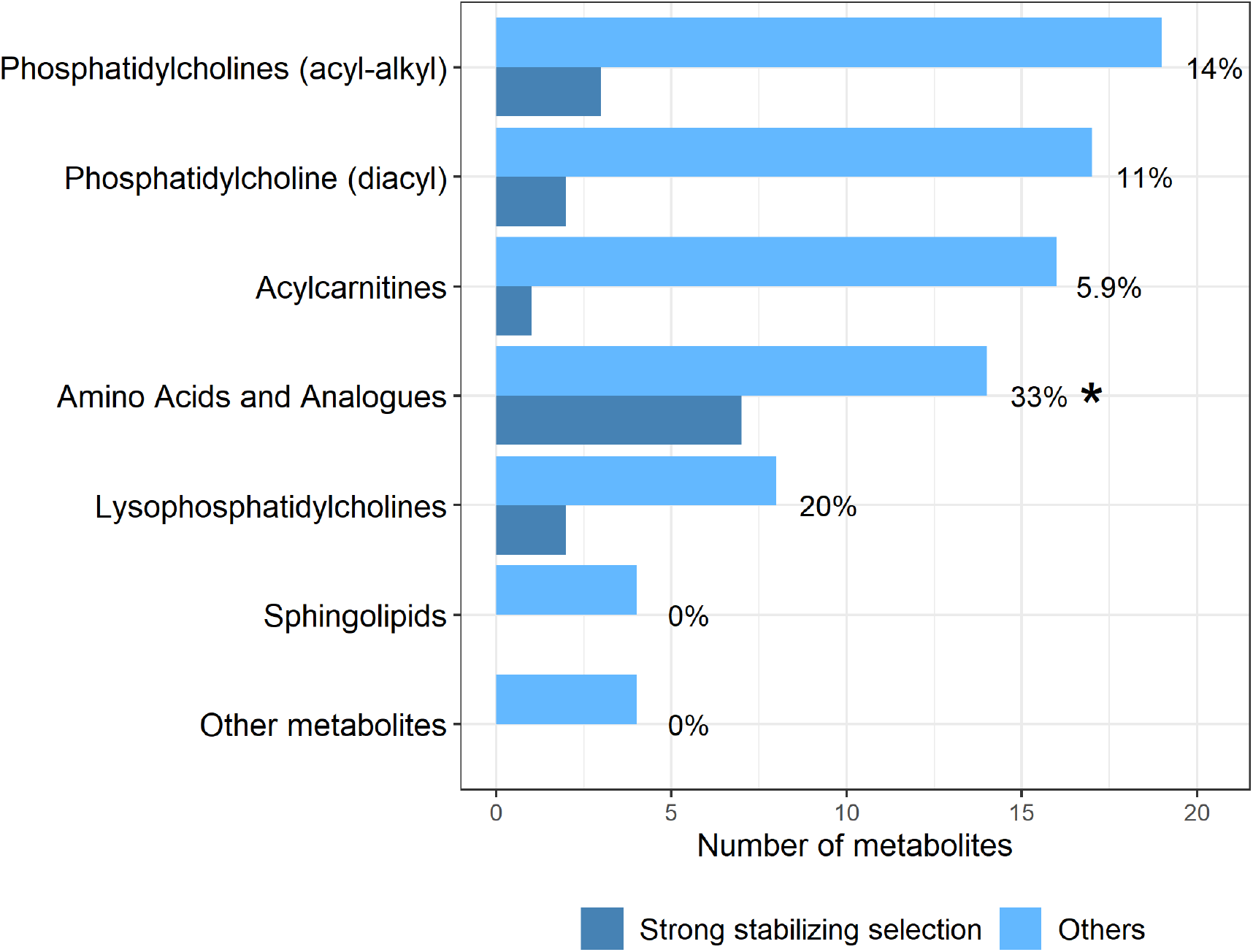
Metabolites under strong stabilizing selection are enriched among amino acids and derivatives. Barplot shows the frequency of metabolites with strong evidence for stabilizing selection (i.e. with selection strength P-value surviving Bonferroni correction) across major metabolite classes. Metabolite classes are based on HMDB. The compound class ‘Amino acids and analogoues’ (i.e. amino acids and their derivatives) are enriched in metabolites with stabilizing selection (P=0.018, two-sided Fisher’s exact test, denoted by asterisk).

### Linking selection strength with the causal effect on cardiometabolic traits

Natural selection to maintain the optimum values of clinically important complex traits might underlie the strong signatures of stabilizing selection on specific metabolites. To test this hypothesis, we first used a Mendelian randomization (MR) approach to unveil the causal links between metabolites and cardiometabolic traits. Applying an inverse variance weighted MR method^15^, we established the causal effects of the 135 metabolites on five complex cardiometabolic traits (Supplementary Table 2): body mass index (BMI), ischaemic heart disease (IHD), systolic blood pressure (SBP), low density lipoproteins (LDL), and type 2 diabetes mellitus (T2D). We identified 30 metabolites with causal effects on LDL with the most significant positive effects on LDL observed for lysoPC a C24:0 (beta = 0.25, P = 1.38 × 10^−35^). Unsurprisingly, some metabolites demonstrated pleiotropic effects on the studied cardiometabolic traits. For example, glutamate was shown to have a negative effect on LDL (beta = -0.39, P = 1.04 × 10^−15^), as well as on IHD (beta = -0.2, P = 2.35 × 10^−5^). In a similar vein, several lipids (PC aa C38:6, SM C16:0, SM C18:0, PC aa C34:1, PC aa C34:4, PC ae C38:1, PC ae C42:2) demonstrated significant positive effects on LDL and weaker positive effects on ischaemic heart disease.

Weak blood pressure-decreasing effects were detected for glycine, phenylalanine, methylglutarylcarnitine, ornithine, and PC ae C36:5, while SBP-increasing was observed for decenoylcarnitine, hexanoylcarnitine, leucine, tryptophan, valine, and SM C18:0. Acylcarnitines (including acetylcarnitine, octadecandienylcarnitine, octadecanoylcarnitine, methylglutarylcarnitine, dodecanoylcarnitine, pimelylcarnitine, tetradecadienylcarnitine, and dodecenoylcarnitine) had modest, but consistent BMI-increasing effect, while phosphatidylcholines and lyso-phosphatiylcholines demonstrated negative causal effects on BMI (Supplementary Table 2). Negative effects on type 2 diabetes (T2D) were detected for PC aa C34:1, PC aa C36:5, ornithine, PC aa C34:3, PC ae C32:1, and PC ae C36:5, while threonine and glutamine had positive causal effects on T2D.

We then interrogated the relationship between the selection strength estimates for the 97 metabolites and their absolute MR effect sizes. Consistent with our hypothesis, we detected negative correlations for all five tested traits, although nominally significant correlation was found only for LDL (R = -0.21, P = 0.04). Thus, metabolites under stronger stabilizing selection show larger effect sizes on these cardiometabolic traits. Note that no significant correlation was observed between the selection strength estimates and the P-values obtained by the MR analysis (Figure S6), which may be due to the fact that, in general, MR P-values strongly depend on the number of genetic markers associated with the risk factors, not only on the causal effect strength.

The most robust negative selection strength estimates with large causal effects were detected for certain phosphatidylcholines (Supplementary Table 2). Overall, these findings indicate that strong stabilizing selection on the plasma concentration of specific phosphatidylcholines may be driven by their causal effect on cardiometabolic health.

### Selection strength correlates with cross-species evolutionary conservation of metabolite levels

The selective forces shaping human metabolism are likely to be shared, at least partly, among mammalian species. If so, metabolites that are under stronger stabilizing selection in human populations are expected to be more evolutionarily conserved in their concentrations over macroevolutionary time scales. To test this hypothesis, we used a recent approach to infer a score that captures the extent of conservation of metabolite concentrations for individual metabolites based on cross-species comparisons^6^. We focused on a metabolomic dataset containing the relative concentrations of 249 metabolites in four major organs of 26 mammalian species, spanning an evolutionary period of ∼200 million ^3^, and calculated an aggregated score of metabolite conservation across the four organs (see Methods). Out of the 97 metabolites with alpha value estimates, 47 were present in the cross-species dataset, including several amino acids, phosphatidylcholines and acylcarnitines (Supplementary Table 2). In line with our expectations, metabolites with lower alpha values (i.e. stronger stabilizing selection) tend to have higher cross-species conservation scores (R=-0.37, p=0.013; Figure 3). This trend indicates that metabolites under stabilizing selection in humans, as detected from GWAS, tend to evolve slowly on macroevolutionary time scales. For example, amino acids frequently show strong negative alpha values and also tend to be highly conserved among mammalian species, while acylcarnitines often display positive alpha values and are among the least conserved metabolites (Figure 3). The presence of this trend is all the more remarkable as cross-species evolution also involves adaptive shifts in metabolite levels outside the human lineage ^3^ that likely diminish the above correlation.

**Figure 3.**
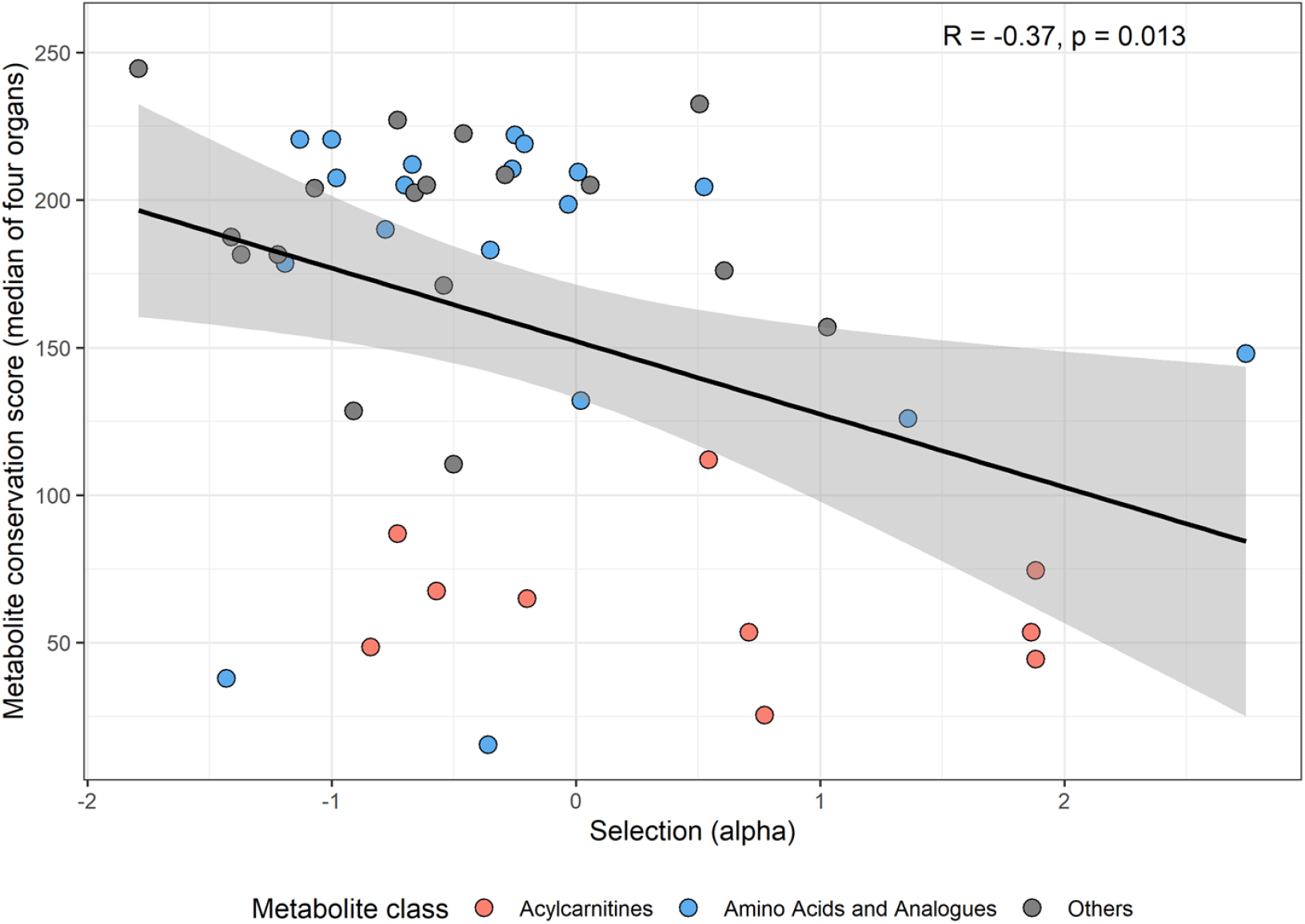
Metabolites under stronger stabilizing selection show stronger evolutionary conservation across mammals. Pearson’s correlation coefficient is indicated on the plot. Evolutionary conservation of metabolite concentrations is estimated by a single ‘metabolite conservation score’ calculated from cross-species metabolomics data in four organs: brain, heart, liver and kidney (see Methods). Line depicts the fitted linear regression. Colours represent two metabolite classes, amino acids and their derivatives (blue) and acylcarnitines (pink).

### Evidence for directional selection on metabolites

While we observed pervasive negative selection for the examined metabolites, these analyses assumed that the current population mean is very close to the optimal concentration value of the given metabolite. If this assumption does not hold, *α*-estimates can be misleading. Such violations can be given rise due to directional selection, whereby not the variance, but the mean value of the metabolite distribution is affected by natural selection. Therefore here we perform an orthogonal analysis to detect directional selection on metabolites based on GWAS data. To investigate the imprint of directional selection on metabolites, we used suggestively (p < 5 × 10^−6^) and genome-wide significantly (p < 5 × 10^−8^) associated SNPs and associated their effect allele frequency with the (signed) effect size of the effect allele. When trait-increasing effects are coupled with elevated effect allele frequency, one may conclude that the optimal trait value is higher than the current population mean. This analysis identified PC aa C40:2 as a metabolite with strong evidence (surviving multiple testing correction at 5%) for positive directional selection (P=4.25 × 10^−5^), i.e. larger effect SNPs tend to have higher allele frequency. This indicates that the population mean value for PC aa C40:2 is below the optimal value for this metabolite and expected to increase in the following generations. Traces of nominally significant directional selection were also detected for arginine (P = 0.028), PC aa C40:3 (P = 0.025), and PC aa C42:5 (P = 0.035) (Figure S4).

## Discussion

We performed the first study, to our knowledge, to assess stabilizing/disruptive and directional selection strength of metabolites using summary statistics from metabolomics GWAS. Leveraging this approach, we were able to estimate stabilizing selection strength for 97 metabolites, via investigating the allele frequency-dependent genetic architecture of metabolomic traits. Information on the strength of selection can be useful in assessing the role of the metabolites as diagnostic, prognostic or treatment response biomarkers.

In order to achieve these results, we first established that the 65-parameter LDscore-weighted BLD-LDAK-Alpha model produces the most robust estimates of selection strength for complex traits, and hence applied it to 135 metabolites with available GWAS summary statistics. Our findings indicate that the majority of the studied amino acids, phosphatidylcholines, lyso phosphatidylcholines, acetylcarnitines, and related compounds with evidence for selection display negative alpha values indicative of stabilizing selection, with only three metabolites showing nominally significant positive selection strength signatures. Our results align with the view that the majority of complex traits are under stabilizing selection that eliminates metabolite-associated genetic variants from the population to avoid deleterious fitness effects ^16^. While it is unsurprising that high-level complex phenotypes, such as reproductive or cardiovascular traits, generally show negative *α* estimates ^8; 9; 11; 17^ due to their close links to fitness, our work expands this notion to molecular traits. Alterations in molecular traits might not necessarily influence higher level phenotypes and fitness, and therefore it has been proposed that such traits are more likely to evolve neutrally ^18^. Our work demonstrates that metabolites, an important layer of molecular traits, also show strong signatures of negative selection in their genetic architectures, albeit less frequently than high-level complex traits ^8^.

When compared to the selection strength profiles of high-level phenotypes identified in our study, the *α* values for the metabolites tend to vary wider and due to the relatively smaller GWAS size available, the estimates are noisier (Figures 1 and S1). Recent technological advancements allowed the detection of hundreds of metabolic compounds that can potentially be used as biomarkers for diseases or drug targets. However, different operating standards and the lack of reference values obtained from healthy subjects lead to large discrepancies in detected metabolite levels across different platforms and laboratories, impeding metabolic profiling ^19^ and downstream analyses, like ours. The MR analysis results demonstrate that many metabolites have causal effect on clinically relevant, complex traits, although, in line with transcriptome-wide MR analyses ^20^, these effects are minor. Still, we hypothesized that clinically more important metabolites would have larger impact on common disease traits. At the same time, if small changes in metabolites lead to disease consequences, their levels are expected to be under stronger stabilizing selection. This prompted us to check whether larger (absolute) causal disease effects couple with stronger (negative) selection values, and we found that selection strength estimates negatively correlated with the absolute causal effect of these metabolites on all tested traits and even nominally significantly so for LDL (R = -0.21, P = 0.04). These observed trends, even if not always strictly statistically significant, support our hypothesis that metabolites with larger impact on clinically important outcomes tend to be under stronger selection. Our results also imply that selection to maintain cardiometabolic health is an important source of evolutionary constraints on certain metabolites.

Strong signatures of stabilizing selection were especially prevalent among amino acids and derivatives. We detected strong negative selection estimates for six amino acids (asparagine, glutamate, glutamine, methionine, proline, tyrosine) in agreement with their essential role in protein synthesis and production of other biologically active substances including thyroid hormones, as well as their involvement in the regulation of the key metabolic pathways necessary for maintenance, growth, reproduction, and immunity. We demonstrated that methionine was characterised by strong negative selection (*α* = -0.98), which may reflect its evolutionary role in initiation of the protein translation. We further obtained the evidence of significant negative selection estimate for proline (*α* = -1.19, P = 3.52 × 10^−23^), which acts as a linchpin of metabolic reprogramming during cancer and related processes, including aging, senescence, and development. LysoPC a C16:1 (*α* = -1.37) was associated with type 2 diabetes, and lysoPC a C20:3 (*α* = -1.79) with blood pressure level ^21; 22^. PC aa C32:1 (*α* = -1.41) along with methionine were identified as one of the key links between homocysteine pathway and telomere length ^23^. Butyrylcarnitine, a short-chain acylcarnitine (C4) and the only member of the acylcarnitine family with negative selection strength estimate (*α* = -0.84), was identified as an informative prognostic marker for neonatal hypoxic-ischemic encephalopathy – condition, characterised by hypoxia triggering a complex response that leads to energy failure, disruption of cellular homeostasis, morphologic changes in microglial cells, and mitochondrial failure ^24^.

Our analyses indicate that certain metabolites might have been shaped by adaptive evolution in the recent evolutionary past of humans. First, we found 3 metabolites (citrulline, lysoPC 20:4, PC ae 40:5) with nominally significant positive alpha estimates. In such cases, the variants associated with the metabolite are under positive selection, potentially indicating disruptive or directional selection on the metabolite concentration ^8^. Directional selection may simply reflect ongoing selection, whereby the population mean has not yet reached the trait optimum. Disruptive selection could emerge due to underdominance ^25^ or could be driven by the pervasive pleiotropy we observe for common variants ^26^. The latter would reflect antagonistic pleiotropy when the same allele (or two alleles in strong linkage disequilibrium) may be beneficial for one trait, but detrimental for another. The strongest positive alpha estimate was detected for citrulline, the key intermediate of the urea cycle, involved also in nitric oxide production ^27^. Clearly, future studies on larger sample sizes should provide further evidence that these metabolites are under directional / disruptive selection and are not simply under relaxed selection. Second, we sought evidence for the action of direction selection using an orthogonal method and found three nominally significant cases as well as a lipid metabolite (PC aa C40:2) with strong statistical evidence for directional selection. In such cases, the population mean is away from the optimum value and is expected to change in the following generations. Intriguingly, several metabolites with signatures of positively selected variants (alpha>0) have been implicated in diseases. Specifically, lysoPC a 20:4 was identified as biomarker for the potentially lifespan-limiting stroke recurrence ^28^, while PC ae 40:5 emerged as a HLA-independent biomarker of multiple sclerosis ^29^. One may speculate that the regulation of such metabolites might be suboptimal in the present environment and hence are more prone to harmful dysregulation during ageing.

We found a remarkable agreement between our estimates of selection based on GWAS data and patterns of metabolite concentration divergence over longer evolutionary time scales. Specifically, we found that metabolites with lower alpha values tend to be more conserved in the concentrations across diverse mammalian species. This finding has at least two general implications. First, it is broadly consistent with the neutral theory of molecular evolution positing that most within-species polymorphisms and between-species divergences at the molecular level are effectively neutral, i.e. permitted rather than favoured by natural selection ^30^. While the theory was originally proposed to explain DNA and protein sequence evolution, it could in principle apply to complex molecular traits as well ^18^. The predominance of neutral evolution of metabolite levels is further supported by our result that metabolites more frequently show signatures of stabilizing selection than those of directional / disruptive selection. Second, the agreement between alpha estimate and between-species conservation score suggests that the selective constraints preserving metabolite levels are at least partly shared between human and other mammalian species, including distantly related taxonomic groups. A recent study suggests that variation in evolutionary conservation across metabolites can be explained by a simple model where natural selection preserves flux through key metabolic reactions while permitting the accumulation of selectively neutral changes in enzyme activities ^6^. Future works should test the extent to which this general model explains stabilizing selection on human metabolite levels.

Last, our results have implications for the understanding of the genetic architecture of molecular traits. While for non-molecular traits no *α* parameter was observed to go below -1, we here report some metabolites showing more extreme selection strength. This threshold has a special meaning, since *α* < -1 indicates that low frequency markers have more per-SNP-heritability than common ones, while *α* > -1 points to an architecture where the average contribution of a common SNP to the trait heritability is more than that of the rare counterparts. This implicates that as opposed to complex traits, for metabolites, the rare variant contribution may be far more important.

Our study has several methodological limitations that can influence interpreting the results. Variable sample size and heritability of the studied trait are key factors determining the power of the study to detect selection strength acting on a trait. Therefore, our analyses cannot be used to establish a priority ranking, but rather to identify some metabolites with significant evidence for selection. Additionally, the directional selection analysis is subject to biases due to confounding by the LD score. Finally, we used the largest available metabolomics GWAS dataset, but the results obtained in our study require further validation using independent study samples and different quantification methods.

## Acknowledgements

This work was supported by the Swiss National Science Foundation (IZSEZ0-194773), the Ministry of Science and Higher Education of Russian Federation Grant No. 075-15-2021-595 (YT), the National Research, Development and Innovation Office ‘Élvonal’ Program KKP 129814 (BP), the “Lendület” program of the Hungarian Academy of Sciences LP2009-013/2012 (BP), the National Laboratory for Health Security, RRF-2.3.1-21-2022-00006 (B.P.) and the European Union’s Horizon 2020 research and innovation program Grant No. 739593 (BP).

## Notes

### Competing Interest Statement

The authors have declared no competing interest.

